# The small GTPase Arf6 functions as a membrane tether in a chemically-defined reconstitution system

**DOI:** 10.1101/2020.11.12.380873

**Authors:** Kana Fujibayashi, Joji Mima

## Abstract

Arf-family small GTPases are essential protein components for membrane trafficking in all eukaryotic endomembrane systems, particularly during the formation of membrane-bound, coat protein complex-coated transport carriers. In addition to their roles in the transport carrier formation, a number of Arf-family GTPases have been reported to physically associate with coiled-coil tethering proteins and multisubunit tethering complexes, which are responsible for membrane tethering, a process of the initial contact between transport carriers and their target subcellular compartments. Nevertheless, whether and how indeed Arf GTPases are involved in the tethering process remain unclear. Here, using a chemically-defined reconstitution approach with purified proteins of two representative Arf isoforms in humans (Arf1, Arf6) and synthetic liposomes for model membranes, we discovered that Arf6 can function as a bona fide membrane tether, directly and physically linking two distinct lipid bilayers even in the absence of any other tethering factors, whereas Arf1 retained little potency to trigger membrane tethering under the current experimental conditions. Arf6-mediated membrane tethering reactions require *trans*-assembly of membrane-anchored Arf6 proteins and can be reversibly controlled by the membrane attachment and detachment cycle of Arf6. The intrinsic membrane tethering activity of Arf6 was further found to be significantly inhibited by the presence of membrane-anchored Arf1, suggesting that the tethering-competent Arf6-Arf6 assembly in *trans* can be prevented by the heterotypic Arf1-Arf6 association in a *cis* configuration. Taken together, these findings lead us to postulate that self-assemblies of Arf-family small GTPases on lipid bilayers contribute to driving and regulating the tethering events of intracellular membrane trafficking.

## Introduction

Small GTPases of the Arf (ADP-ribosylation factor) family, belonging to the Ras superfamily, are known to be essential protein components for intracellular membrane trafficking in all eukaryotic endomembrane systems and, in general, to function through cooperation with their specific interactors, termed Arf effectors (D’Souza-Schorey and Chavrier, 2006; Donaldson and Jackson, 2011; Jackson and Bouvet, 2014; Sztul et al., 2019). In particular, a large body of prior studies on Arf-family small GTPases have established their pivotal functions during the formation of membrane-bound, coat protein complex-coated transport carriers (e.g., secretory and endocytic transport vesicles) at the donor membranes of subcellular compartments (D’Souza-Schorey and Chavrier, 2006; Sztul et al., 2019): Arf1 recruits the COPI (coat protein complex I) subunits for initiating vesicle formation in the retrograde Golgi-to-endoplasmic reticulum (ER) pathway (Spang et al., 1998; Bremser et al., 1999); Arf6 binds to the AP-2 adaptor complex in the clathrin-mediated endocytic pathway (Paleotti et al., 2005); and Sar1p assembles with Sec23/24p and Sec13/31p to form the COPII (coat protein complex II) coat and thereby promote cargo sorting and vesicle budding in the anterograde ER-Golgi trafficking pathway (Matsuoka et al., 1998; Sato and Nakano, 2004; Sato and Nakano, 2005). In addition to their roles in the transport carrier formation above, it should also be noted that a number of Arf-family small GTPases, including Arl (Arf-like) GTPases, have been reported to physically interact with various long coiled-coil tethering proteins and multisubunit tethering complexes as a non-coat Arf effector (Donaldson and Jackson, 2011; Sztul et al., 2019), which include golgin GMAP-210 for Arf1 (Drin et al., 2008), the exocyst complex for Arf6 (Prigent et al., 2003), Golgin-97 and Golgin-245 for Arl1 (Lu and Hong, 2003), the GARP complex for Arl5 (Rosa-Ferreira et al., 2015), and the HOPS complex for Arl8 (Khatter et al., 2015). All of these tethering proteins or tethering complexes are thought to be directly involved in membrane tethering, vesicle tethering, or vesicle capture, a process of the initial physical contact of membrane-bound transport carriers with their target subcellular membrane compartments (Waters and Pfeffer, 1999; Yu and Hughson, 2010; Gillingham and Munro, 2019). This tethering process is vital for determining the spatiotemporal specificity of membrane trafficking (Waters and Pfeffer, 1999; Yu and Hughson, 2010; Gillingham and Munro, 2019), before the following SNARE-mediated membrane docking and fusion steps, another critical layers for conferring the fidelity of membrane trafficking (McNew et al., 2000; Parlati et al., 2002; Furukawa and Mima, 2014). Nevertheless, Arf-family small GTPases have not been recognized as the so-called tethering factors (Waters and Pfeffer, 1999; Yu and Hughson, 2010; Gillingham and Munro, 2019), and thus, whether and how indeed Arf-family proteins contribute to the tethering process remain unclear. In this study, to thoroughly explore the molecular functions of Arf-family proteins in membrane tethering, we have employed reconstituted membrane tethering assays with Arf-anchored proteoliposomes, which were prepared from purified human Arf1 and Arf6 proteins and synthetic liposomes bearing defined lipid species. Using the chemically-defined reconstitution approach, here we report that Arf6, but not Arf1, can directly and physically tether two distinct lipid bilayers in the absence of any other tethering factors or Arf effectors.

## Materials and Methods

### Protein expression and purification

Bacterial expression vectors for human Arf1 (UniProtKB: P84077) and Arf6 (UniProtKB: P62330) were constructed using a pET-41 Ek/LIC vector kit (Novagen), as described for protein expression of human Rab-family small GTPases (Tamura and Mima, 2014; Inoshita and Mima, 2017; Segawa et al., 2019; Ueda et al., 2020). DNA fragments encoding these two Arf-family proteins with the additional sequences for a human rhinovirus (HRV) 3C protease-cleavage site and a His12-tag at the N-terminus were amplified by PCR using KOD-Plus-Neo polymerase (Toyobo) and Human Universal QUICK-Clone cDNA II (Clontech) as a template cDNA and then cloned into a pET-41 Ek/LIC vector (Novagen). The GST-His6-tagged forms of His12-Arf1 and His12-Arf6 proteins were expressed in *E. coli* BL(DE3) (Novagen) cells harboring the pET-41-based expression vectors constructed. After inducing expression in the presence of IPTG (final 0.3 mM) at 25°C for 4 h, cells were harvested by centrifugation, resuspended in RB150 (20 mM Hepes-NaOH, pH 7.4, 150 mM NaCl, 10% glycerol) containing 5 mM MgCl_2_, 1 mM DTT, 1 mM PMSF, and 1 μg/ml pepstatin A, freeze-thawed in liquid nitrogen and a water bath at 30°C, lysed by sonication, and ultracentrifuged with a 70 Ti rotor (Beckman Coulter; 50,000 rpm, 75 min, 4°C). The supernatants were mixed with COSMOGEL GST-Accept beads (Nacalai Tesque) and incubated with agitation (4°C, 5 h), followed by washing the protein-bound beads in RB150 with 5 mM MgCl_2_ and 1 mM DTT. The washed beads were resuspended in the same buffer containing HRV 3C protease (8 units/ml in final; Novagen) and incubated without agitation (4°C, 16 h) to cleave off and elute His12-Arf proteins from the beads. His12-Arf1 and His12-Arf6 proteins eluted were harvested by centrifugation (15,300 × *g*, 10 min, 4°C). Human Rab5a-His12 was purified as described (Segawa et al., 2019; Ueda et al., 2020). Protein concentrations of purified His12-Arf1, His12-Arf6, and Rab5a-His12 were determined using Protein Assay CBB Solution (Nacalai Tesque) and BSA for a standard protein. Nucleotide loading of human Arf-family small GTPases was performed by incubating purified Arf proteins in the presence of GTP or GDP (1 mM), EDTA (2 mM), and MgCl_2_ (1 mM), as described previously (Antonny et al., 1997; Drin et al., 2008).

### GTPase activity assay

GTP hydrolysis activities of purified His12-Arf proteins were assayed by quantitating free phosphate molecules released in the hydrolytic reactions, using the Malachite Green-based reagent Biomol Green (Enzo Life Sciences) as described for the activity assays with Rab-family small GTPases (Inoshita and Mima, 2017; Segawa et al., 2019). Purified His12-Arf1 and His12-Arf6 proteins (2 μM final) were mixed with GTP or GTPγS (1 mM final) in RB150 containing 5 mM MgCl_2_ and 1 mM DTT (100 μl each), incubated (30°C, 1 h), and then supplemented with the Malachite Green reagent (100 μl each). After incubation with the reagent (30°C, 30 min), the absorbance at 620 nm (A620) of the reactions (200 μl each) was measured with a DU720 spectrophotometer (Beckman Coulter). The raw A620 data were corrected by subtracting the A620 values of the control reactions without His12-Arf proteins. Concentrations of phosphate molecules released in the reactions were calculated using the corrected A620 data and the A620 values of phosphate standards (Enzo Life Sciences). Means and standard deviations of the specific GTPase activities (μM phosphate/min/μM protein) were determined from three independent experiments.

### Liposome preparation

All of the non-fluorescent lipids for liposome preparation, including POPC (1-palmitoyl-2-oleoyl-phosphatidylcholine), POPE (1-palmitoyl-2-oleoyl-phosphatidylethanolamine), liver PI (phosphatidylinositol), POPS (1-palmitoyl-2-oleoyl-phosphatidylserine), cholesterol, and DOGS-NTA (1,2-dioleoyl-sn-glycero-3-{[N-(5-amino-1-carboxypentyl) iminodiacetic acid]-succinyl}), were from Avanti Polar Lipids. The two fluorescence-labeled lipids, Rh-PE (rhodamine-PE) and FL-PE (fluorescein-PE), were from Molecular Probes. Lipid mixes used were prepared in chloroform with the lipid compositions of 41% (mol/mol) POPC, 17% POPE, 10% liver PI, 5% POPS, 20% cholesterol, 6% DOGS-NTA, and 1% Rh-PE or FL-PE, dried up by evaporating chloroform with a stream of nitrogen gas, and subsequently resuspended in RB150 containing 5 mM MgCl_2_ and 1 mM DTT (final 8 mM lipids) by vortexing vigorously and incubating with agitation (37°C, 1 h). After freeze-thawing in liquid nitrogen and a water bath at 30°C, lipid suspensions were extruded 25 times through polycarbonate filters (pore diameters, 200 nm; Avanti Polar Lipids) in a mini-extruder (Avanti Polar Lipids) preheated at 40°C. The liposome solutions prepared were stored at 4°C and used within a week for liposomes turbidity assays and fluorescence microscopy.

### Liposome turbidity assay

The intrinsic membrane tethering activities of human Arf-family small GTPases were assayed by monitoring turbidity changes of liposome solutions in the presence of purified Arf proteins, as described for the turbidity assays with human Rab-family small GTPases (Tamura and Mima, 2014; Inoshita and Mima, 2017; Mima, 2018; Segawa et al., 2019; Ueda et al., 2020) and Atg8-family proteins (Taniguchi et al., 2020). Purified His12-Arf1, His12-Arf6, or Rab5a-His12 (0.5 – 3 μM final) and DOGS-NTA-bearing liposomes (200-nm diameter; final 1 mM total lipids), which had been separately preincubated (30°C, 10 min), were mixed in RB150 containing 5 mM MgCl_2_ and 1 mM DTT (160 μl each), applied to a 10-mm path-length cell (105.201-QS, Hellma Analytics), and immediately subjected to monitoring the turbidity changes with measuring the optical density changes at 400 nm (ΔOD400) for 5min with 10-sec intervals in a DU720 spectrophotometer at room temperature. The ΔOD400 values obtained from the kinetic turbidity assays were further quantitatively analyzed by curve fitting using ImageJ2 (National Institutes of Health) and the logistic function formula, *y* = *a*/(1 + *b* × exp(-*c* × *x*)), in which *y* is the ΔOD400 value and *x* is the time (min), as described (Segawa et al., 2019; Taniguchi et al., 2020; Ueda et al., 2020). The maximum tethering capacities of Arf and Rab proteins were defined as the maximum ΔOD400 values of the fitted sigmoidal curves at *t* = ∞, and they can be calculated as “*a*” from the logistic formula. The initial tethering velocities were defined as the maximum slopes of the fitted curves, and they can be calculated as “*c* × *a*/4” from the logistic formula. Means and standard deviations of the maximum capacities (ΔOD400) and the initial velocities (ΔOD400/min) were determined from three independent experiments. All of the kinetic turbidity data shown were obtained from one experiment and typical of those from three independent experiments.

### Fluorescence microscopy

Fluorescence microscopy assays of Arf-mediated membrane tethering were employed using a LUNA-FL fluorescence cell counter (Logos Biosystems) and LUNA cell-counting slides (L12001, Logos Biosystems), as described (Segawa et al., 2019; Taniguchi et al., 2020; Ueda et al., 2020). Purified His12-Arf1 or His12-Arf6 (1 or 2 μM final) and extruded liposomes bearing DOGS-NTA and fluorescence-labeled Rh-PE or FL-PE lipids (200-nm diameter; final 2 mM lipids) were preincubated separately (30°C, 10 min), mixed in RB150 containing 5 mM MgCl_2_ and 1 mM DTT (80 μl each), and further incubated (30°C, 1 h). The liposome suspensions incubated with Arf proteins were applied to the cell-counting slides (15 μl/well) and subjected to microscopic assays using the LUNA-FL fluorescence cell counter to obtain the bright field images and rhodamine- and fluorescein-fluorescence images. Particle sizes of liposome clusters in the rhodamine-fluorescence images were analyzed using ImageJ2 with setting the lower intensity threshold level to 150, the upper intensity threshold level to 255, and the minimum particle size to 10 pixels^2^, which corresponds to approximately 10 μm^2^.

## Results and Discussion

Among twenty-nine members of the Arf small GTPase family in humans, including five Arf GTPases, twenty Arl GTPases, and two Sar1 GTPases (Sztul et al., 2019), in the current reconstitution studies on membrane tethering, we selected the two representative Arf isoforms, Arf1 and Arf6, both of which have been well-known to play key roles in regulating the secretory and endocytic trafficking pathways (D’Souza-Schorey and Chavrier, 2006; Sztul et al., 2019) and interact with putative membrane tethers or tethering factors such as coiled-coil tethering proteins and multisubunit tethering complexes (Prigent et al., 2003; Drin et al., 2008; Donaldson and Jackson, 2011). These two Arf-family proteins (Arf1 and Arf6), which share over 65% sequence identity (**Figure 1A**), and most of the other Arf isoforms are small monomeric globular proteins (20-25 kDa) consisting of the amphipathic helices at the N-terminus (around 10 residues; the helix α1 in **Figure 1A**) and the following conserved Ras-superfamily GTPase domains (G-domains; 160-170 residues; **Figure 1B**), and typically, they are further post-translationally modified by a myristoyl group at the N-terminus, which functions as a lipid anchor required for their stable attachment on the membrane surface (Donaldson and Jackson, 2011; Sztul et al., 2019). To mimic the native membrane-bound state of myristoylated Arf proteins in the present chemically-defined reconstitution systems, we prepared recombinant Arf1 and Arf6 proteins as the N-terminal polyhistidine-tagged forms (His12-Arf proteins; **Figure 1B, C**) that can be efficiently and specifically bound to DOGS-NTA lipids incorporated into synthetic liposomal membranes used. Purified His12-Arf1 and His12-Arf6 proteins (**Figure 1C**) retained their intrinsic GTP-hydrolysis activities, specifically converting GTP to GDP and a free phosphate (**Figure 1D**), which were comparable to those of Rab5a-His12 used as the control of a tethering-active small GTPase (**Figure 2**) and the other human Rab-family proteins tested in our prior studies on reconstituted membrane tethering (Inoshita and Mima, 2017; Segawa et al., 2019). Thus, these results confirmed that His12-Arf1 and His12-Arf6 in the current preparations were correctly folded and a functional protein with a native-like active G-domain.

**Figure 1.**
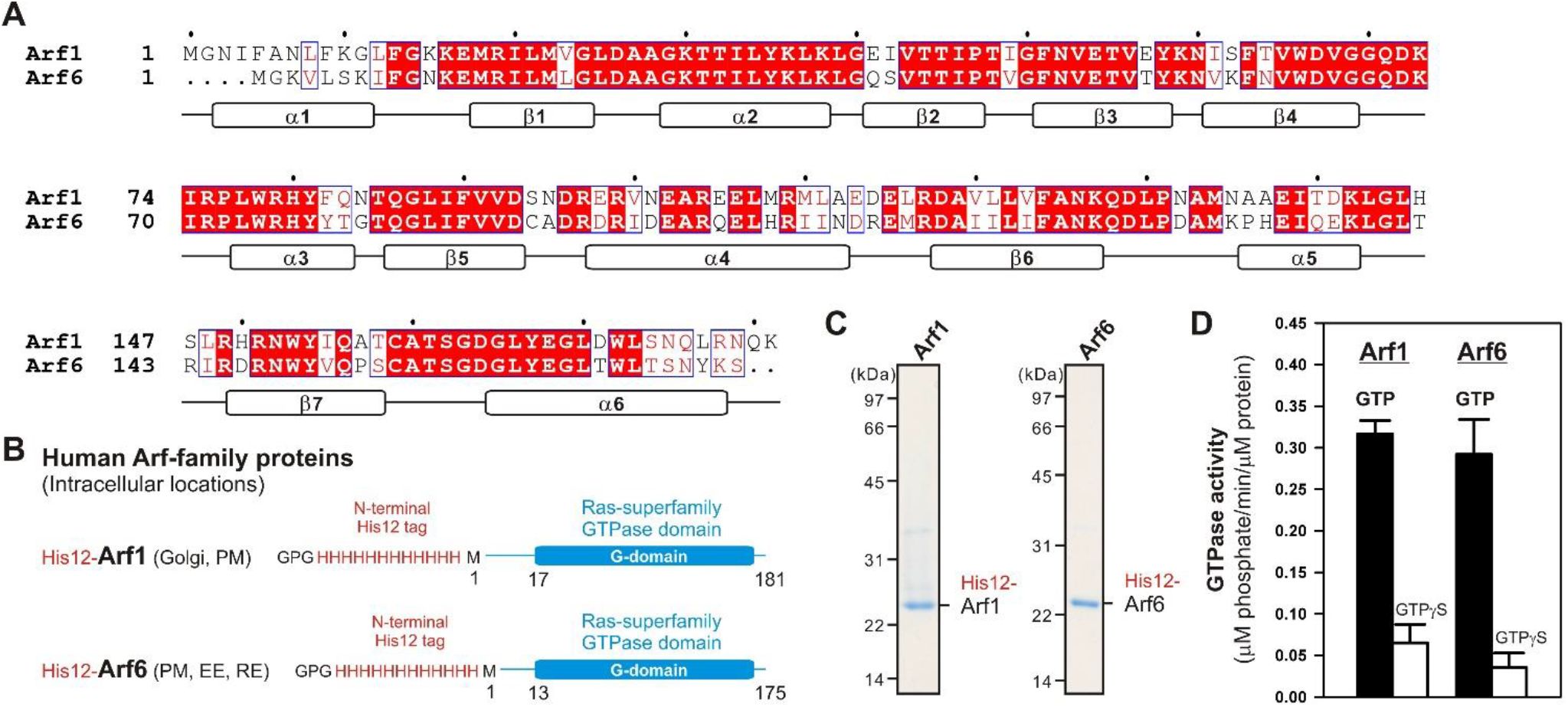
Human Arf-family small GTPases used in the current reconstitution studies. (**A**) Sequence alignment of human Arf1 and Arf6 proteins. Amino acid sequences of these two Arf isoforms were obtained from UniProtKB (https://www.uniprot.org/), aligned using ClustalW (https://www.genome.jp/tools-bin/clustalw), and rendered by ESPript 3.0 (http://espript.ibcp.fr/ESPript/ESPript/). Identical and similar amino acid residues in the sequence alignment are highlighted in red boxes and in red characters, respectively. Secondary structures determined by the crystal structure of human Arf1 containing the residues 2-181 (PDB code, 1HUR), including six α-helices (α1-α6) and seven β-strands (β1-β7), are indicated at the bottom of the alignment. (**B**) Schematic representation of recombinant N-terminally His12-tagged human Arf1 (His12-Arf1) and Arf6 (His12-Arf6) proteins used in the current studies, which contain three extra residues (Gly-Pro-Gly) at the extreme N-terminus, an N-terminal His12-tag, and the Met1 – Lys181 residues for Arf1 or the Met1 – Ser175 residues for Arf6. The intracellular locations of Arf1 and Arf6 indicated include the Golgi apparatus (Golgi), plasma membrane (PM), early endosome (EE), and recycling endosome (RE). (**C**) Coomassie blue-stained gels of purified His12-Arf1 and His12-Arf6 proteins, which were tested in the reconstituted liposome tethering assays in **Figures 2-4**. (**D**) GTP hydrolysis activities of purified recombinant His12-Arf1 and His12-Arf6 small GTPases. Purified Arf proteins (2 μM final) were incubated with 1 mM GTP or GTPγS (30°C, 1 h) in RB150 containing 5 mM MgCl_2_ and 1 mM DTT and then assayed for the specific GTPase activities by measuring the free phosphate molecules released in the hydrolytic reactions.

**Figure 2.**
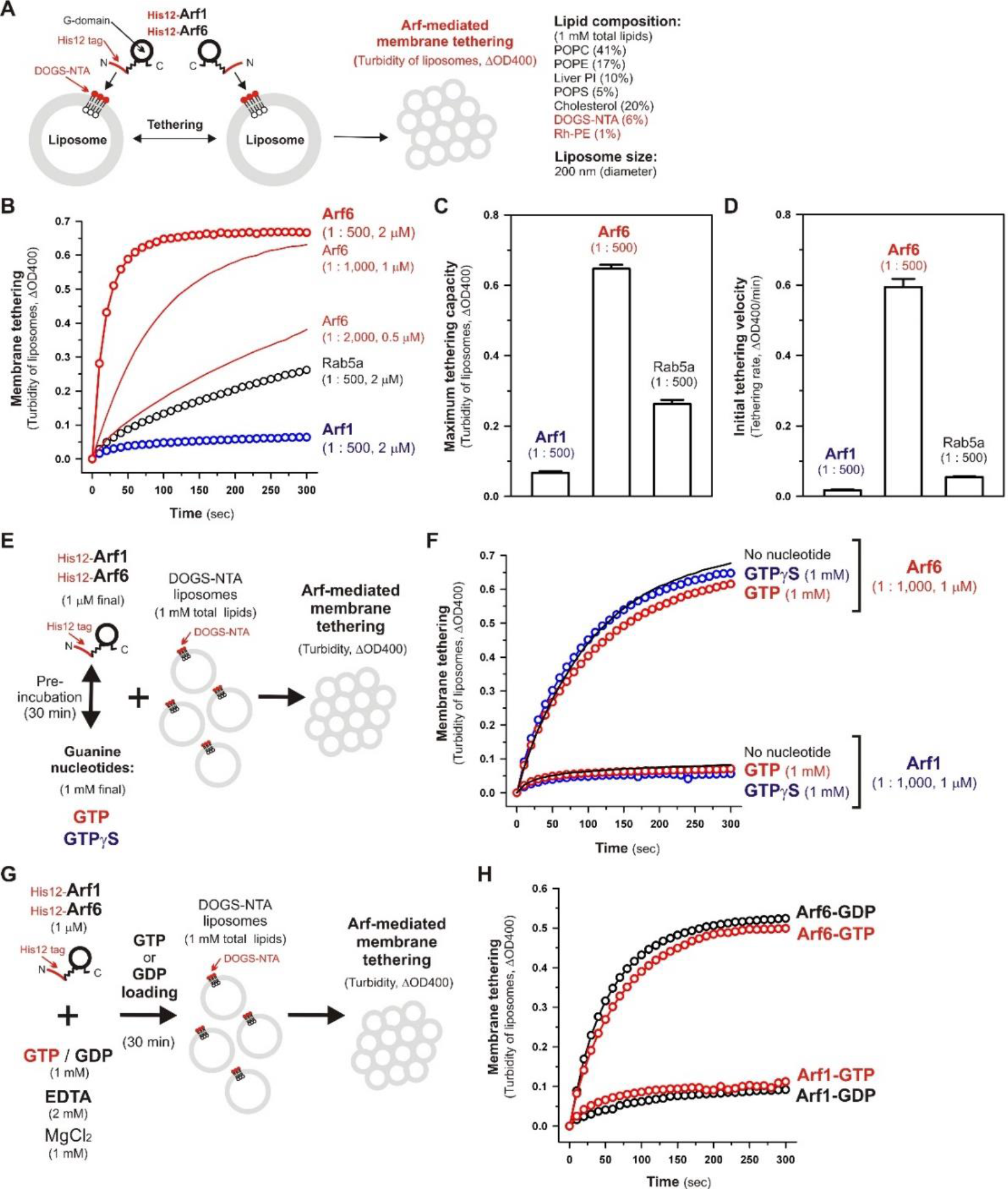
Human Arf6, but not Arf1, can directly mediate membrane tethering in a chemically-defined reconstitution system. (**A**) Schematic representation of liposome turbidity assays for testing the intrinsic membrane tethering potency of human Arf-family proteins. (**B-D**) The intrinsic membrane tethering activities of human Arf1 and Arf6 proteins in kinetic liposome turbidity assays. Purified His12-Arf1, His12-Arf6, and Rab5a-His12 for the control of a tethering-competent small GTPase (0.5, 1, or 2 μM in final) were mixed with DOGS-NTA-bearing liposomes (200-nm diameter; 1 mM total lipids in final), immediately followed by assaying the turbidity changes by measuring the optical density changes at 400 nm (ΔOD400) for 5min with 10-sec intervals (**B**). Maximum tethering capacities (ΔOD400) (**C**) and initial tethering velocities (ΔOD400/min) (**D**) of the turbidity reactions at 2 μM His12-Arf1, His12-Arf6, and Rab5a-His12 proteins were determined using a sigmoidal curve fitting method. (**E**) Schematic representation of liposomes turbidity assays with His12-Arf proteins in the presence of guanine nucleotides. (**F**) Addition of guanine nucleotides, GTP and GTPγS, has little or no effect on the intrinsic tethering activities of Arf1 and Arf6. After preincubating His12-Arf1 and His12-Arf6 proteins (1 μM final) with GTP or GTPγS (1 mM final) or without any guanine nucleotides (30°C, 30 min), kinetic liposome turbidity assays were employed as in (**B**). (**G**) Schematic representation of liposomes turbidity assays with GTP-loaded or GDP-loaded Arf proteins. (**H**) Nucleotide-loaded state has little or no effect on the tethering activities of Arf1 and Arf6. After incubating His12-Arf1 and His12-Arf6 proteins (1 μM) with GTP or GDP (1 mM), EDTA (2 mM), and MgCl_2_ (1 mM) at 30°C for 30 min, kinetic liposome turbidity assays for the nucleotide-loaded forms of Arf1 (Arf1-GTP, Arf1-GDP) and Arf6 (Arf6-GTP, Arf6-GDP) were employed as in (**B**). The protein-to-lipid molar ratios used, ranging from 1:500 to 1:2,000 (mol/mol), are indicated (**B, C, D, F**). Error bars, SD.

Using purified His12-Arf1 and His12-Arf6 proteins (**Figure 1**) and extruded 200-nm protein-free liposomes bearing DOGS-NTA lipids and the complex lipid composition (**Figure 2A**), which roughly mimics the compositions of subcellular membrane compartments in mammalian cells (van Meer et al., 2008; Vance, 2015; Yang et al., 2018), we first examined whether these two Arf isoforms can exhibit their intrinsic potency to directly and physically tether two distinct lipid bilayers by employing kinetic liposome turbidity assays (**Figure 2B-D**), as established for Rab-mediated membrane tethering in our earlier studies (Tamura and Mima, 2014; Inoshita and Mima, 2017; Mima, 2018; Segawa et al., 2019; Ueda et al., 2020). Strikingly, the kinetic turbidity assays revealed that Arf6 triggered rapid and efficient tethering of liposomes when tested at the Arf protein concentrations ranging from 0.5 to 2 μM, which correspond to the Arf-to-lipid molar ratios (mol/mol) of 1:2,000 to 1:500 (red open circles and red solid lines, **Figure 2B**). It should be noted that these Arf-to-lipid molar ratios are similar to the protein-to-lipid ratios tested for evaluating the tethering activities of putative membrane tethers or tethering factors in prior reconstitution experiments, which include the typical ratios of 1:400 for golgin GMAP-210 (Drin et al., 2008), 1:330 for Vps21p (Lo et al., 2012), 1:2,000 for HOPS (Ho and Stroupe, 2015; Ho and Stroupe, 2016), 1:100-1:5,000 for Rab-family small GTPases (Segawa et al., 2019; Ueda et al., 2020), 1:800 for Atg8p (Nair et al., 2011), and 1:100-1:5,000 for LC3B and GATE-16 (Taniguchi et al., 2020). By contrast, the other Arf isoform, Arf1, exhibited little or no tethering activity under the current experimental conditions tested at 2 μM Arf proteins with the 200-nm liposomes (blue open circles, **Figure 2B**), consistent with our prior results showing no significant tethering activity of Arf1 with larger 400-nm liposomes (Segawa et al., 2019). To further evaluate the novel tethering activity of Arf6 found here, we also tested the non-Arf, Rab-family small GTPase, Rab5a, which has been recognized as the most tethering-active Rab isoform (Segawa et al., 2019), in the same kinetic turbidity assays and compared the tethering activities of Arf6 and Rab5a (**Figure 2B**). Although Rab5a retained its decent tethering activity for the 200-nm liposomes (black open circles, **Figure 2B**), Arf6 exhibited about 2.5-fold higher tethering capacity and over 10-fold higher initial tethering velocity than those values of Rab5a (**Figure 2C, D**). These results established that Arf6 can function as a more potent membrane tether than the highly-active Rab-family tether, Rab5a, for the 200-nm liposomes at least.

Since Ras-superfamily small GTPases, including Arf and Rab GTPases, are generally thought to function in the active GTP-bound forms on membranes (Rojas et al., 2012), we next examined the intrinsic tethering activities of Arf1 and Arf6 in the presence of guanine nucleotides (**Figure 2E, F**). His12-Arf1 and His12-Arf6 were pre-incubated with GTP or the non-hydrolyzable analog GTPγS under the conditions similar to those used in the GTPase activity assays in **Figure 1D** (final 1 μM Arf proteins and 1 mM guanine nucleotides, 30°C, 30 min; **Figure 2E**) and then subjected to the kinetic turbidity assays as in **Figure 2B** (**Figure 2F**). With respect to both of the Arf isoforms, Arf1 and Arf6, the presence of either GTP or GTPγS had little or no effect on their tethering capacities and initial tethering velocities in the turbidity assays (**Figure 2F**), establishing that Arf-mediated membrane tethering is achieved in a guanine nucleotide-independent manner. This consistent with our earlier experiments for reconstituted Rab-mediated tethering, in which the tethering activities of human Rab-family proteins were totally insensitive to the addition of guanine nucleotides (Tamura and Mima, 2014). In addition, nucleotide-independent Arf-mediated tethering was further demonstrated by reconstituted tethering assays with nucleotide-loaded Arf proteins, in which GTP-loaded Arf6 (Arf6-GTP) and Arf1 (Arf1-GTP) proteins had tethering capacities and kinetics almost identical to those of their GDP-loaded forms (**Figure 2G, H**). Considering that the Rab11a effectors, class V myosins, can specifically stimulate membrane tethering mediated by the cognate Rab11a in a GTP-dependent manner (Inoshita and Mima, 2017), perhaps specific Arf effectors can confer the nucleotide-dependence of Arf-driven tethering reactions.

Although the intrinsic tethering potency of human Arf small GTPases was not able to be controlled by the nucleotide-bound states in the current reconstitution systems using His12-tagged Arf proteins and DOGS-NTA-bearing liposomes (**Figure 2E-H**), it should be noted that, in living cells, native myristoylated Arf-family proteins are soluble and inactive in the GDP-bound state at the cytoplasm, but they can be functionally active in the GTP-bound state on subcellular membranes through inserting their N-terminal myristoyl lipid anchors and amphipathic helices to membrane surfaces (Antonny et al., 1997; Liu et al., 2009; Liu et al., 2010). Liposome co-sedimentation assays for recombinant Arf6 proteins with or without a His12 tag indicated that their stable membrane association was not achieved by the amphipathic helices alone even in the GTP-loaded states (**Figure S1**). Thus, this suggests that the artificial membrane anchoring via an N-terminal His12 tag and a DOGS-NTA lipid can bypass the GTP requirement for membrane binding of Arf proteins and their following tethering functions in a reconstitution system.

To further strengthen the experimental evidence for the intrinsic membrane tethering potency of the Arf-family GTPase, Arf6, we next performed fluorescence microscopic imaging assays to analyze particle sizes of liposome clusters induced by Arf6-mediated tethering (**Figure 3**). When incubated Arf proteins with fluorescence-labeled 200-nm liposomes bearing Rh-PE (Arf-to-lipid molar ratios of 1:1,000, 30°C, 1 h; **Figure 3A**), Arf6 induced efficiently the formation of massive liposome clusters, yielding the total particle area of 410,000 ± 38,000 μm^2^ and the average particle size of 1,300 ± 530 μm^2^, whereas Arf1 exhibited little potency to induce large liposome clusters that were detectable by the current imaging assay (**Figure 3B, C**). In addition, Arf6-induced liposome clusters were able to be dissociated into undetectable small particles by incubating in the presence of imidazole (250 mM) that blocks the association of His12-Arf6 with a DOGS-NTA lipid on liposomes (**Figure 3D, E**). This establishes that Arf6-mediated tethering is a non-fusogenic, reversible tethering reaction, and it can be strictly controlled by the membrane attachment and detachment cycle of Arf6 on lipid bilayers. Using the same microscopic imaging assay but with two types of the fluorescence-labeled liposomes bearing DOGS-NTA and either Rh-PE or FL-PE (**Figure 3F-H**), we also asked whether Arf6-mediated membrane tethering requires *trans*-assembly between membrane-anchored Arf6 proteins on two opposing membranes to be tethered, as established for reconstituted membrane tethering reactions mediated by human Rab-family small GTPases (Tamura and Mima, 2014; Inoshita and Mima, 2017; Segawa et al., 2019; Ueda et al., 2020) and by Atg8-family proteins (Taniguchi et al., 2020). When DOGS-NTA lipids were present in both of the Rh-bearing and FL-bearing liposomes (**Figure 3F**), Arf6 retained the tethering capacity to induce massive clusters containing the two fluorescence-labeled liposomes, yielding the total particle are of 430,000 ± 23,000 μm^2^ (**Figure 3F**). However, by omitting a DOGS-NTA lipid from the FL-labeled liposomes (**Figure 3G**), Arf6 lost its ability to tether them, forming only Rh-labeled large liposome clusters (the total particle are of 180,000 ± 13,000 μm^2^; middle panel, **Figure 3G**). Moreover, as expected, the tethering activity of Arf6 was completely abrogated when DOGS-NTA lipids were not present in either of the two fluorescence-labeled liposomes (**Figure 3H**). The requirement of membrane-anchored Arf6 proteins for membrane tethering was further established by liposome turbidity assays in the presence of soluble Arf6 proteins lacking an N-terminal His12 tag, showing that a vast molar excess of untagged Arf6 had no potency to competitively inhibit membrane tethering mediated by membrane-anchored Arf6 proteins (**Figure S2**). Therefore, these results demonstrate that Arf6-mediated tethering is selectively driven by *trans*-assembly of the membrane-anchored form of Arf6 proteins between two apposing lipid bilayers.

**Figure 3.**
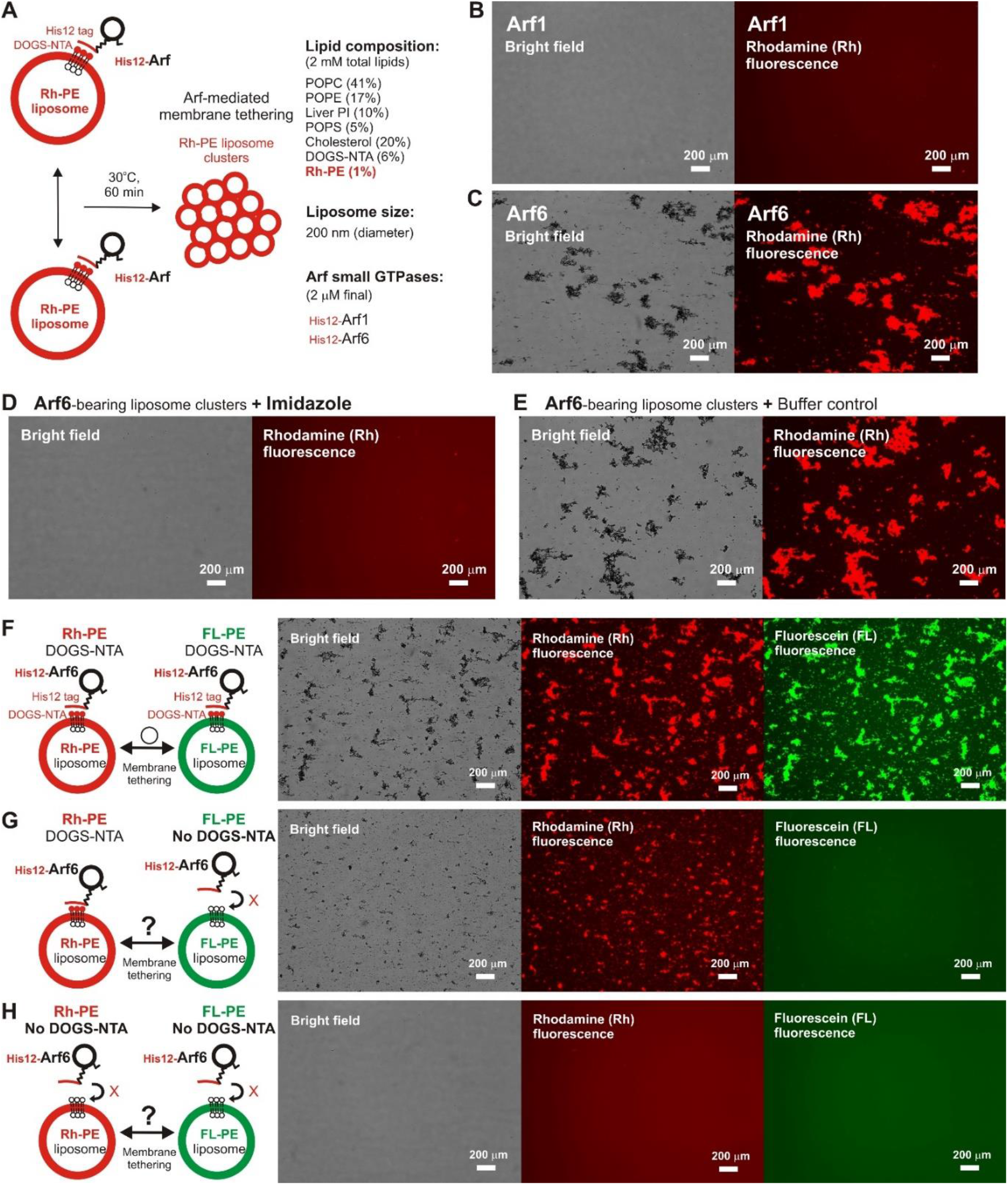
Fluorescence microscopic analysis of human Arf6-mediated membrane tethering. (**A**) Schematic representation of fluorescence microscopy-based tethering assays with human Arf-family small GTPases. (**B, C**) Purified His12-Arf1 (**B**) and His12-Arf6 (**C**) proteins (final 2 μM) were mixed with fluorescence-labeled liposomes bearing Rh-PE and DOGS-NTA (200-nm diameter; final 2 mM lipids), incubated (30°C, 1 h), and subjected to fluorescence microscopy. (**D, E**) Reversibility of Arf6-mediated membrane tethering. Purified His12-Arf6 was incubated with Rh-PE/DOGS-NTA-bearing liposomes as in (**C**) (30°C, 1 h), supplemented with imidazole (final 250 mM) which causes the dissociation of His12-Arf6 from the DOGS-NTA-containing liposomes (**D**) or with the buffer control (**E**), further incubated (30°C, 1 h), and subjected to fluorescence microscopy. (**F-H**) Requirement of *trans*-assembly between membrane-anchored Arf proteins for Arf6-mediated membrane tethering. Purified His12-Arf6 (final 2 μM) was incubated with two types of fluorescence-labeled liposomes, Rh-PE-bearing liposomes and FL-PE-bearing liposomes (200-nm diameter; final 1 mM lipids for each), as in (**C**) (30°C, 1 h) and subjected to fluorescence microscopy. DOGS-NTA lipids were present in both of the fluorescence-labeled liposomes (**F**), only in the Rh-PE liposomes (**G**), or not present in either of the liposomes (**H**). Scale bars, 200 μm.

We previously reported that membrane tethering mediated by Rab-family small GTPases can be triggered not only by “homotypic” Rab-Rab assembly in *trans* (e.g., Rab5a-Rab5a, Rab6a-Rab6a, and Rab7a-Rab7a; Tamura and Mima, 2014; Inoshita and Mima, 2017; Segawa et al., 2019), but also by “heterotypic” *trans*-assembly between two different Rab isoforms, including Rab1a-Rab6a, Rab1a-Rab9a, and Rab1a-Rab33b (Segawa et al., 2019). Therefore, although Arf1 had little tethering potency in a homotypic fashion (**Figures 2B, 3B**), next we employed reconstituted tethering assays in the presence of both Arf1 and Arf6, to test whether the inactive isoform Arf1 can be functional in the tethering reactions through interacting with Arf6 on membranes (**Figure 4A**). Nevertheless, in the kinetic turbidity assays (**Figure 4B**), the tethering activity of Arf6 was substantially diminished by adding a 2-fold molar excess of Arf1 (red open circles, **Figure 4B**), exhibiting an approximately 6-8-fold decrease in the maximum tethering capacity (**Figure 4C**) and in the initial tethering velocity (**Figure 4D**), whereas the addition of extra Arf6, instead of Arf1, significantly enhanced the rate of Arf6-mediated liposome tethering (black solid line, **Figure 4B**). This specific inhibitory effect on Arf6-mediated tethering requires the membrane-anchored form of Arf1, as the tethering activity of Arf6 was unable to be inhibited by either soluble untagged Arf1 or a His12 peptide (**Figure S3**). Fluorescence microscopy assays also demonstrated that the formation of massive liposome clusters induced by Arf6-mediated tethering was totally inhibited by the presence of membrane-anchored Arf1 on liposomes (**Figure 4E, F**). Thus, these results suggest that Arf1 can interact with Arf6 on lipid bilayers, not in *trans* for initiating the tethering events of two distinct membranes, but in a *cis* configuration, thereby preventing the self-assembly of Arf6 proteins in *trans*. We further investigated guanine nucleotide dependence of the heterotypic *cis*-interactions between Arf6 and Arf1 on lipid bilayers by employing turbidity assays with the GTP/GDP-loaded forms of Arf6/Arf1 proteins (**Figure 4G, H**). Arf1-GTP retained the potency to substantially inhibit membrane tethering mediated by Arf6-GTP (red open circles, **Figure 4G**), consistent with the prior results obtained in the absence of guanine nucleotides (**Figure 4B**). Interestingly, the GDP-loaded form of Arf1 exhibited relatively moderate inhibitory effect on Arf6-GDP-mediated tethering (black open circles, **Figure 4G**). This may reflect that *cis*- and *trans*-assemblies of Arf-family proteins on membranes are driven by distinct modes of the protein-protein interactions.

**Figure 4.**
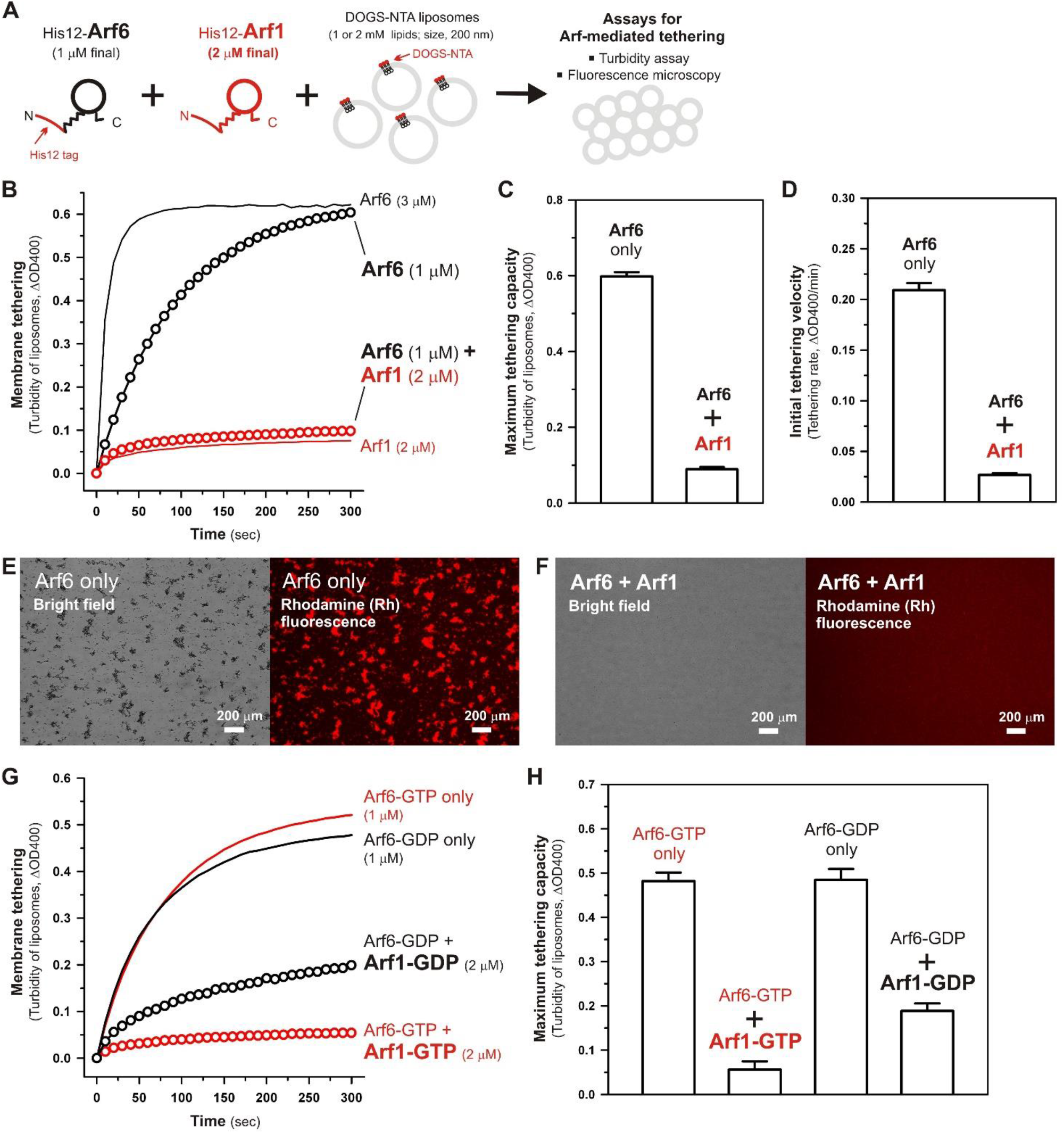
Arf-mediated membrane tethering in the presence of the heterotypic pair of Arf6 and Arf1. (**A**) Schematic representation of reconstituted liposome tethering assays for testing membrane tethering mediated by the heterotypic combination of Arf6 and Arf1. (**B-D**) Kinetic liposome turbidity assays. Purified His12-Arf6 (final 1 or 3 μM), His12-Arf1 (final 2 μM), or both (1 μM Arf6 and 2 μM Arf1 in final) were mixed with DOGS-NTA-bearing liposomes (200-nm diameter; final 1 mM lipids) and immediately assayed for the turbidity changes (**B**), as in **Figure 2B**. Maximum tethering capacities (**C**) and initial tethering velocities (**D**) were determined for the turbidity reactions with Arf6 only (final 1 μM) and the reactions with both Arf6 and Arf1, as in **Figure 2C, D**. (**E, F**) Fluorescence microscopy. Purified His12-Arf6 (final 1 μM) was incubated (30°C, 10 min) in the absence (**E**) or the presence (**F**) of purified His12-Arf1 (final 2 μM), mixed with Rh-PE/DOGS-NTA-bearing liposomes (200-nm diameter; final 2 mM lipids), further incubated (30°C, 1 h), and subjected to fluorescence microscopy, as in **Figure 3B, C**. (**G, H**) Kinetic liposome turbidity assays for nucleotide-loaded Arf proteins. GTP-loaded or GDP-loaded Arf6 (Arf6-GTP or Arf6-GDP; 1 μM) was incubated with Arf1-GTP or Arf1-GDP (2 μM), mixed with DOGS-NTA-bearing liposomes, and assayed for the turbidity changes (**G**), as in (**B**). Maximum tethering capacities (**H**) were determined for the turbidity reactions, as in (**C**). Scale bars, 200 μm.

Taken together, using a chemically-defined reconstitution approach with purified proteins of Arf1 and Arf6 in humans and synthetic liposomal membranes, we uncovered that Arf6 can directly mediate reversible membrane tethering reactions through the self-assembly in *trans* between two opposing lipid bilayers (**Figures 2-3**). Despite the fact that Arf6 functions as a bona fide membrane tether in the reconstituted system, the other isoform Arf1, which shares over 65% sequence identity with Arf6 and also retains the GTP hydrolysis activity comparable to that of Arf6 (**Figure 1A, D**), exhibited little or no tethering activity under the same experimental conditions (**Figures 2-3**). In addition, the current tethering assays in the presence of both these two Arf isoforms revealed that membrane-anchored Arf1 can significantly inhibit the tethering activity of Arf6 (**Figure 4**), reflecting the heterotypic Arf1-Arf6-assembly in *cis* on lipid bilayers. These findings on the novel molecular functions of Arf6 and Arf1 in membrane tethering lead us to postulate that self-assemblies of Arf-family small GTPases in *trans* and in *cis* can directly contribute to driving and regulating the tethering events of membrane trafficking in eukaryotic cells, perhaps through collaborating with their specific effectors including coiled-coil tethering proteins and multisubunit tethering complexes. Future studies will need to be focused on mechanistic details of Arf-mediated membrane tethering, particularly determining the dimer interface required for *trans*-assembly of Arf molecules on lipid bilayers. Recent experimental and computational works on the membrane-associated dimers of K-Ras and H-Ras small GTPases have proposed that their putative dimer interfaces are located at the C-terminal allosteric lobes in the G-domains, not within the N-terminal effector lobes which contain the switch I, switch II, and inter-switch regions (Muratcioglu et al., 2015; Prakash et al., 2017; Spencer-Smith et al., 2017; Abankwa and Gorfe, 2020). Although it remains totally unknown whether the dimer interfaces of K-Ras and H-Ras are conserved through all of the members belonging to the Ras superfamily, we speculate that the C-terminal allosteric lobes of Arf small GTPases, as well as Rab-family GTPases, are involved in their *trans*-assemblies on membranes, as reconstituted membrane tethering reactions mediated by Arf and Rab proteins are found to be completely insensitive to the guanine nucleotide-bound states.

## Author contributions

JM designed the research and wrote the manuscript. JM and KF performed the experiments and analyzed the data.

## Acknowledgements

We thank Megumi Shinguu (Institute for Protein Research, Osaka University) for the contribution to preparing purified proteins of human Arfs and Rab5a. This study was in part supported by the Grants-in-Aid for Scientific Research from the Ministry of Education, Culture, Sports, Science and Technology, Japan (MEXT) (to JM).

## Supplemental Figures

**Figure S1.**
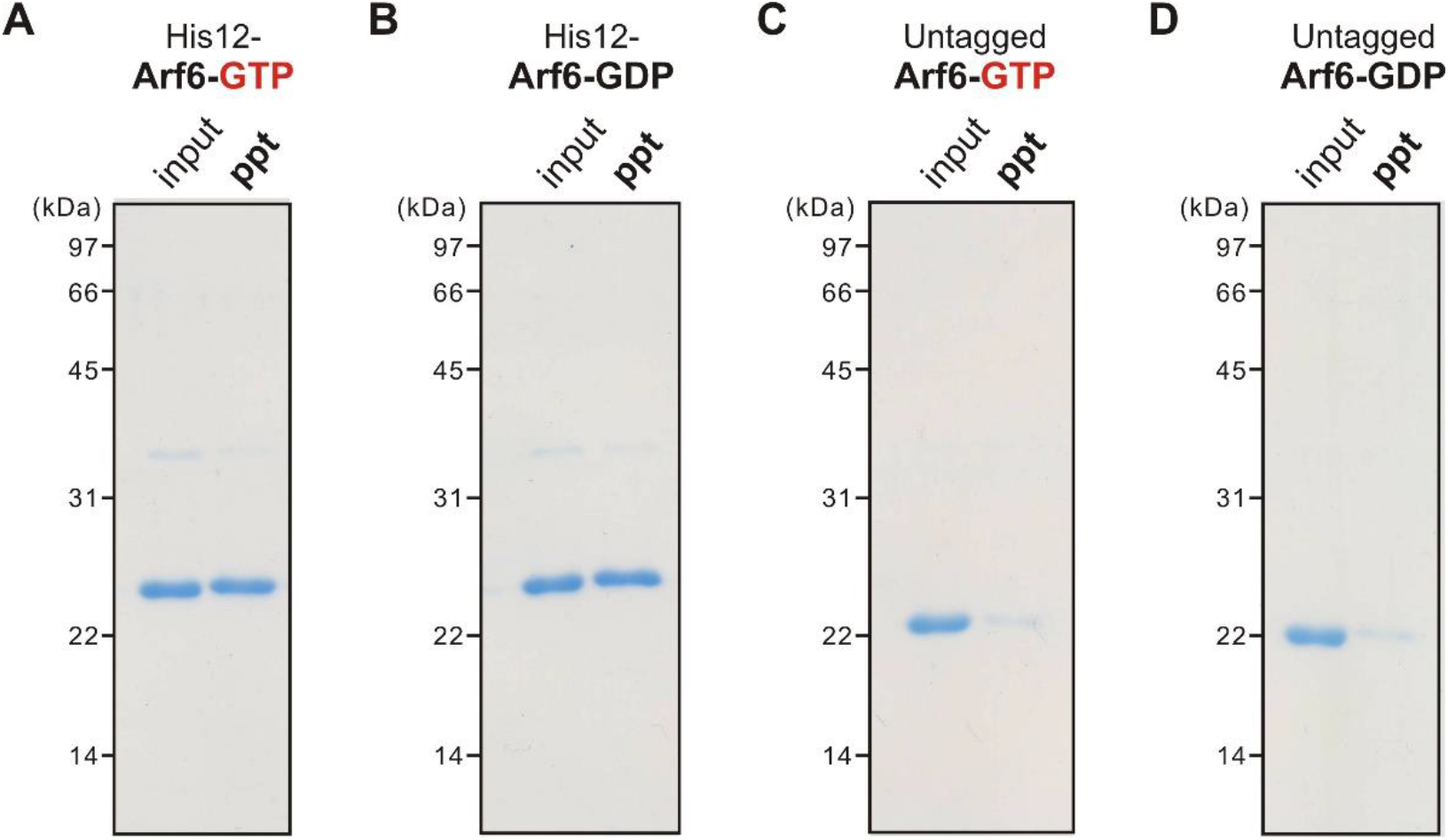
Liposome co-sedimentation assays testing the membrane association of nucleotide-loaded Arf6 proteins. (**A-D**) Purified nucleotide-loaded Arf6 proteins (4 μM), including His12-Arf6-GTP (**A**), His12-Arf6-GDP (**B**), untagged Arf6-GTP lacking a His12 tag (**C**), and untagged Arf6-GDP (**D**), were incubated with DOGS-NTA-bearing liposomes (2 mM lipids; 1,000-nm diameter) in RB150 containing 5 mM MgCl_2_ and 1 mM DTT (30°C, 30 min), centrifuged (20,000 × *g*, 30 min, 4°C), and analyzed by SDS-PAGE and CBB staining for precipitates (*ppt*) obtained after centrifugation.

**Figure S2.**
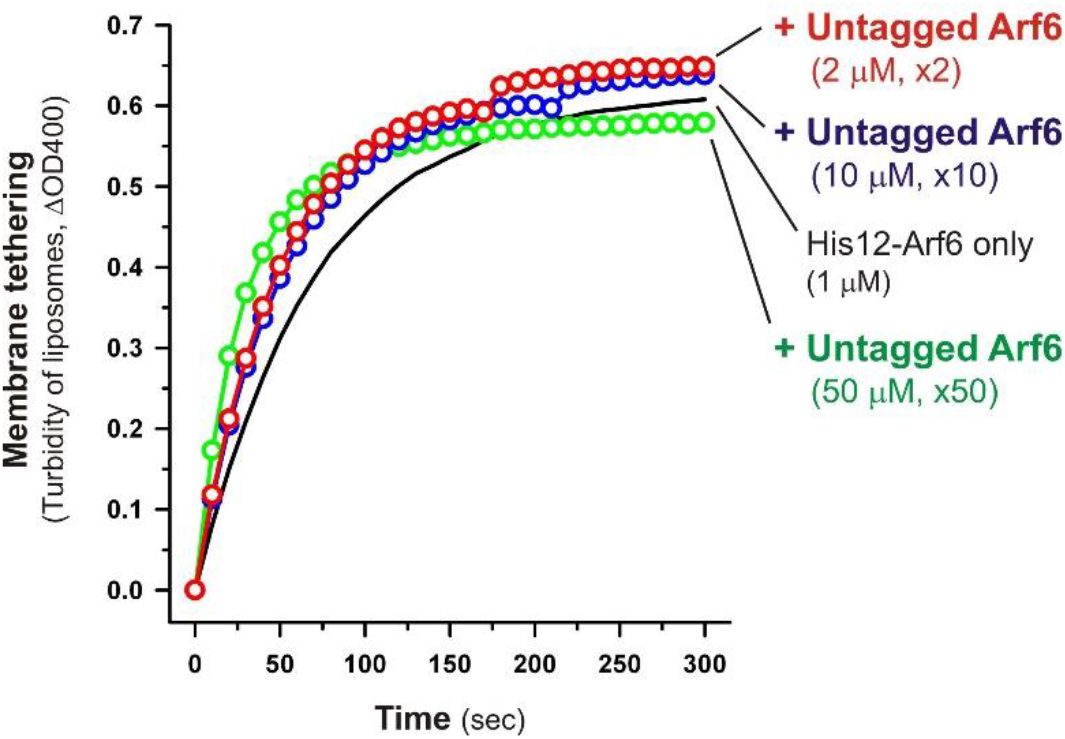
Selective *trans*-assembly between membrane-anchored Arf6 proteins in reconstituted membrane tethering. Purified His12-Arf6 proteins (1 μM) were mixed with excess amounts of untagged Arf6 proteins lacking a His12-tag membrane anchor (2 μM, 10 μM, or 50 μM), incubated (30°C, 10 min), supplemented with DOGS-NTA-bearing liposomes (1 mM lipids; 200-nm diameter), and subjected to liposome turbidity assays, as in **Figure 2B**.

**Figure S3.**
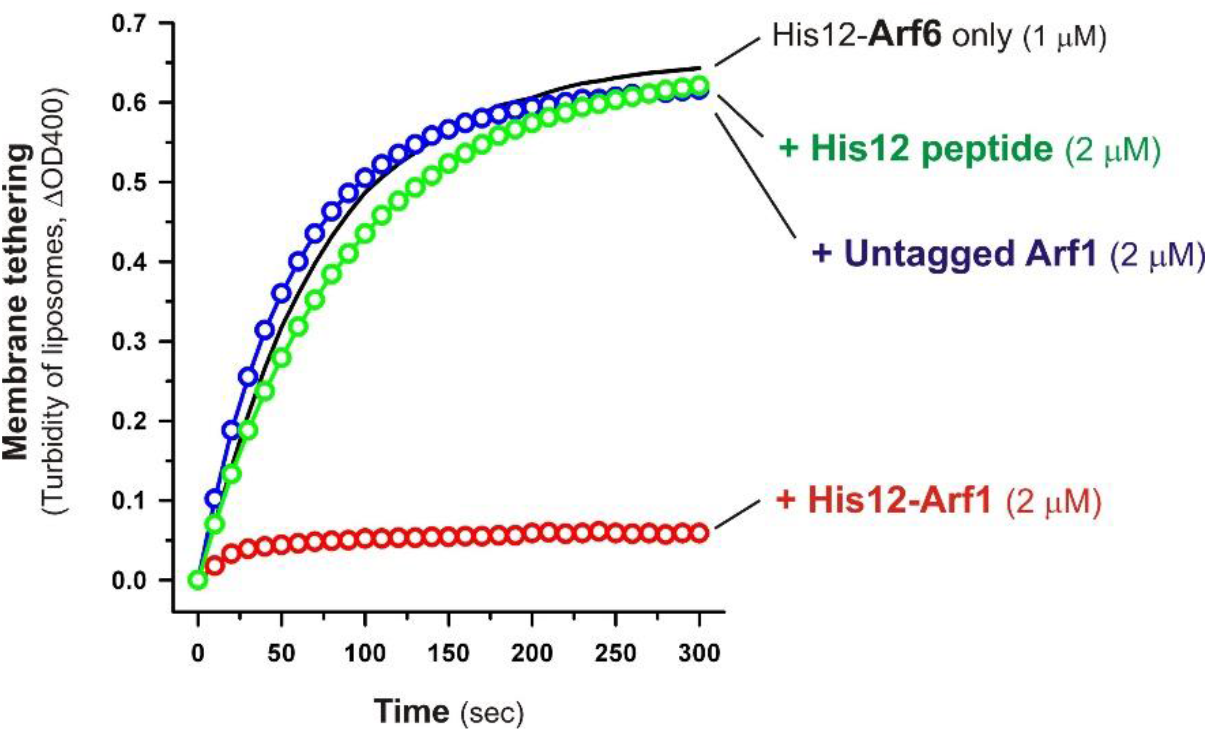
Specific inhibitory effect of membrane-anchored Arf1 on Arf6-mediated membrane tethering. His12-Arf6 (1 μM) was mixed with His6-Arf1 (2 μM), untagged Arf1 lacking a His12 tag (2 μM), or a His12 peptide (2 μM), incubated (30°C, 10 min), mixed with DOGS-NTA-bearing liposomes (1 mM lipids; 200-nm diameter), and subjected to liposome turbidity assays, as in **Figure 4B**.

